# HCVTyper: A Nextflow pipeline for hepatitis C virus genome assembly, genotyping and antiviral resistance detection

**DOI:** 10.1101/2025.10.21.683612

**Authors:** Jon Bråte, Kamilla Heddeland Instefjord, Kristian Alfsnes, Kathrine Stene-Johansen

## Abstract

**Motivation:** High-throughput sequencing is increasingly used for the characterization of hepatitis C virus (HCV), enabling genotyping, detection of mixed infections, and identification of resistance-associated variants (RAVs). However, the bioinformatics analysis of capture-based or metagenomic HCV datasets remains challenging due to high genome diversity, co-infections, and the need for reproducible workflows.

**Results:** We present *HCVTyper*, a reproducible bioinformatics pipeline for HCV genome analysis implemented in Nextflow. *HCVTyper* integrates quality control, taxonomic classification, two-step reference-based mapping, consensus generation, *de novo* assembly, and antiviral resistance annotation. Benchmarking with simulated mixtures and real-world datasets demonstrated robust identification of both major and minor genotypes across a wide abundance range, with user-adjustable thresholds for co-infection detection. Integration with HCV-GLUE enables sensitive detection of RAVs from mapping files, including low-frequency variants. The pipeline is fully containerized (Docker) and produces comprehensive reports to facilitate interpretation.

**Availability and implementation:** *HCVTyper* is open-source and available at: https://github.com/folkehelseinstituttet/hcvtyper

## Introduction

Hepatitis C virus (HCV) strains are now routinely characterized by high-throughput genome sequencing, which enables simultaneous determination of genotype, detection of mixed infections, and identification of antiviral resistance-associated variants (RAVs). This has created a demand for robust and comprehensive bioinformatics workflows that not only provide clinically relevant interpretations but are also scalable, reproducible, and portable across different computing environments.

HCV is an enveloped virus carrying a positive-sense, single-stranded RNA genome of ∼9.6 kb (Chevaliez and Pawlotsky 2006). The genome encodes a single polyprotein that is post-translationally processed into structural and non-structural protein. HCV exhibits extensive genetic diversity: current classification by the International Committee on Taxonomy of Viruses (Walker et al. 2020) recognize eight major genotypes and more than 90 subtypes, with diversity within a single subtype reaching up to ∼15% (Smith et al. 2014).

Highly effective direct-acting antivirals (DAAs) against HCV are available, which target the NS3 protease, NS5A and NS5B polymerase (Brzdek et al. 2023). Although HCV treatment increasingly relies on newer, pan-genotypic DAAs, it has not eliminated the need for genotype-guided therapy for certain populations such as DAA-experienced patients with treatment failures (Brzdek et al. 2023; Pawlotsky et al. 2020). However, because of the high mutation rate and high genetic variation of HCV, RAVs frequently arise naturally or selected during treatment (Howe et al. 2022; Wyles and Luetkemeyer 2017), underscoring the need for sensitive detection methods.

This high genetic diversity poses a challenge to genotyping and RAV identification. Within a host, HCV exists as a dynamic population of closely related variants of the same genotype (“quasispecies”) (Domingo et al. 2021; Kao et al. 1995; Martell et al. 1992). In addition to this intra-genotype diversity, mixed infections with distinct genotypes (co-infections) are well documented (Buckton et al. 2006; Cunningham et al. 2015; McNaughton et al. 2018). In such cases, one strain typically dominates (“major strain”), while others are present at lower abundance (“minor strains”). Detecting both reliably is challenging: reference-based approaches may miss minor strains if reads are mapped only to a single reference sequence, while *de novo* assembly can in principle capture all variants but often produces fragmented assemblies that are difficult to interpret. The problem is further complicated by naturally occurring recombinant HCV genomes (Hedskog et al. 2015; Raghwani et al. 2012), which reference-based methods may misclassify if an appropriate recombinant reference is not present in the database used for classification. In practice, a combination of reference-mapping and *de novo* assembly is often the most robust strategy.

A second challenge arises from the quasispecies nature of HCV populations. Approaches that collapse reads into a single consensus sequence, whether reference-based or *de novo*, will inevitably fail to capture low-frequency variants within each genotype. This is critical because RAVs may exist as clinically relevant variants within the quasispecies (Caputo et al. 2020; Pawlotsky 2016; Perales et al. 2018). Sensitive detection therefore requires analysis of the underlying read depth rather than consensus sequences alone.

Therefore, we need bioinformatics methods for HCV that can fully exploit whole-genome high-throughput sequence data for in-depth strain characterization and resistance identification. Next-generation sequencing (NGS) can in principle sequence the entire HCV viral population from a single sample, and resolve mixed infections (Qiu et al. 2015). Capture-based NGS yields near-complete HCV genomes, even at low viral loads and can accurately resolve mixed-genotype infections (Thomson et al. 2016).

Several general viral genome assembly pipelines exist, such as *viralrecon* (Patel et al. 2023) and *viralmetagenome* (Klaps et al. 2025), and some HCV-specific tools such as *MiCall* (Lapointe et al. 2021) support assembly or genotyping. To the best of our knowledge, no openly available, HCV-tailored workflow integrates genome assembly, genotyping (including co-infections), and DAA resistance analysis from whole-genome NGS data.

To address this gap, we developed *HCVTyper*, a portable pipeline built in Nextflow (Di Tommaso et al. 2017) and following the nf-core best practices (P. A. Ewels et al. 2020). The pipeline is explicitly genotype-agnostic: HCV reads are identified and mapped to a broad set of references covering the entire HCV diversity, then remapped to candidate major and minor strain references before consensus building. DAA resistance analysis is performed on the mapping files using HCV-GLUE (Singer et al. 2018), which allows for the detection of low-abundant RAVs directly from the mapped NGS reads. *HCVTyper* is particularly suited for capture-based Illumina datasets and for samples with co-infections or divergent genotypes. All tools are containerized using Docker, ensuring reproducibility without local installation.

## Implementation

*HCVTyper* is implemented in Nextflow (Di Tommaso et al. 2017) using the nf-core toolkit (P. A. Ewels et al. 2020) to produce the pipeline base template files. All software dependencies are packaged as Docker containers to increase portability and reproducibility and reduce the need for manual installation of software (it is also possible to use Singularity or Conda as environment managers, however currently this will not allow for DAA identification with HCV-GLUE or read-mapping with the Tanoti mapper (Vattipally 2017). The pipeline supports the simultaneous analysis of multiple samples via a samplesheet with sample IDs and FASTQ file paths. Nextflow enables efficient parallelization of the different samples and processes, making it ideal for high-throughput analyses. It also allows for deployment both on local and HPC/cloud systems.

A typical workflow comprises the following steps:

1. Pre-processing:
  - Produce raw sequence data statistics using FastQC (Andrews 2010).
  - Removal of sequencing adapters and low-quality bases using either Fastp (Chen et al. 2018) or Cutadapt (Martin 2011).
  - Taxonomic classification of reads running Kraken2 (Wood et al. 2019) against the PlusPFP-8 database (B. Langmead).
  - Identification of HCV reads by running Kraken2 against a provided HCV-specific database.
2. Read mapping and assembly:
  - Identification of potential major and minor strains by mapping HCV-classified reads with Bowtie2 (Ben Langmead and Salzberg 2012) to a provided set of HCV-reference sequences.
  - Re-mapping the HCV-classified reads against only the candidate major and minor reference sequences using Bowtie2.
  - Create consensus sequences from the mapping files using iVar (Grubaugh et al. 2019).
  - *De novo* assembly and genotyping of HCV-classified reads into contigs using Spades (Bankevich et al. 2012).
3. Antiviral resistance detection:
  - Identification of antiviral resistance using HCV-GLUE (Singer et al. 2018).
4. Reporting:
  - Final outputs include consensus FASTA(s) for each sample, genotyping report with detailed mapping statistics, antiviral resistance summary, and a MultiQC (P. Ewels et al. 2016) report aggregating logs, QC and software versions.

An example command to run the pipeline (with Docker) might be:

~~~
nextflow run folkehelseinstituttet/HCVTyper -r v1.0.6 \
--input samplesheet.csv \
--outdir <OUTDIR> \
-profile docker
~~~

where samplesheet.csv lists each sample and the paths to its paired FASTQ files. A minimal test mode (-profile test) is provided with example data. Additional parameters (e.g. resource limits, path to custom databases and reference sequence) can be set via command-line flags as documented. The pipeline’s source code, documentation and issue tracker are on GitHub (https://github.com/folkehelseinstituttet/hcvtyper).

### Co-infections (major and minor strains)

To perform genotyping, and identify major and minor strains, the pipeline first maps all HCV-classified reads against the complete set of HCV reference sequences. The reference sequence with the largest number of mapped reads is then selected as the candidate major strain, provided that at least 500 reads are mapped (default, can be adjusted with the *--minRead* parameter) and at least 30% of the reference is covered at a depth of ≥5 reads (default, can be adjusted with the *--minCov* parameter). The genotype and subtype information stored in the reference header is used to call major genotype and subtype.

To identify a potential co-infection (minor strain), the pipeline selects the reference with the highest coverage that belongs to a genotype different from the identified major strain. The only exception is the distinction between genotype 1a and 1b, which are treated as sufficiently divergent to allow co-infection calls between them. The same thresholds for read depth and coverage are applied to the candidate minor strains.

Note that there is a recombinant strain between subtypes 2k and 1b present in the database. If this is detected, the pipeline will not allow for a co-infection with either genotypes 1 or 2.

### Pipeline verification

#### Simulated data

We simulated paired-end Illumina MiSeq reads from different HCV reference genomes using InSilicoSeq v2.0.1 (Gourlé et al. 2018). We simulated three datasets; two mixed datasets (named sim1 and sim2) representing a co-infection of genotypes 1a (based on reference HQ850279 downloaded as fasta format from NCBI GenBank) and 1b (reference AJ238799) and a co-infection of genotypes 2a (reference NC_009823) and 3a (reference NC_009824), and a third dataset (sim3) with only a single genotype 4a present (reference NC_009825).

A total of 1 million read pairs were simulated for each dataset. For the co-infections we let one genotype occupy 70% of the reads (i.e., “major” strain) and the other genotype occupy 30% of the reads (“minor” strain). For all genotypes in all datasets we inserted a RAV at low abundance (20% of the reads per genotype). For genotype 1a we inserted the RAV Q30R by changing nucleotide 6196 from A to G. For genotype 1b we inserted L31M by changing nucleotide 6347 from T to A. For genotype 2a L/M31M was inserted by changing nucleotide 6338 from A to G and for genotype 3a M28K was inserted by changing nucleotide 6356 from T to A. For genotype 4a L30A was inserted by changing two nucleotides, position 6238 from T to G and 6284 from T to C. The simulated datasets are available for download from figshare at 10.6084/m9.figshare.30405847.

The coverage distribution along each genome was simulated using the “--lognormal” option in InSilicoSeq. Reads were simulated for each genotype separately and then combined to obtain the specified abundance ratios. An example command is: *iss generate --genomes GtA*.*fa --model MiSeq --cpus 6 –output gt1a_sim --coverage lognormal --n_reads 140000*

For all three samples the pipeline correctly identified the simulated genotypes and the low-frequent RAVs (Table 1). For the co-infected samples, the number of HCV-classified reads that mapped to the “major” and “minor” strains roughly corresponded to their expected abundance ratios based on the number of simulated reads. However, note that as all HCV-classified reads are mapped to both strains, some reads will map to conserved areas in both reference genomes and hence these numbers differ slightly.

**Table 1:** Pipeline validation on simulated datasets.” Sample” refers to the simulated datasets described in the text. “Theoretical genotypes” shows the expected genotypes and their relative abundances as indicated in “Theoretical ratio”. “Identified genotypes” are the genotypes identified by *HCVTyper* (major:minor). “Mapped reads ratio” shows the relative proportions of reads mapped to the major and minor genotypes. The number of mapped reads are shown in “Mapped reads”. These are counted after removing duplicate reads and the numbers are rounded to the nearest thousand (“k”). “Coverage (%)” represents the percent of reference nucleotides covered by at least 10 mapped reads. “Differences to reference” indicates the number of single-nucleotide differences between the *HCVTyper* consensus sequence and the reference used for simulation. “RAV detected” shows whether the low-frequency resistance-associated variant (RAV) inserted into the simulated data was detected.

| Sample | Theoretical genotypes | Theoretical ratio | Identified genotypes | Mapped reads ratio | Mapped reads | Coverage (%) | Differences to reference | RAV detected |
| --- | --- | --- | --- | --- | --- | --- | --- | --- |
| sim1 | 1a:1b | 7:3 | 1a:1b | 6.5:3 | 647k:298k | 100:99.9 | 4:1 | Yes:Yes |
| sim2 | 2a:3a | 7:3 | 2a:3a | 6.2:3 | 629k:303k | 100:99.9 | 0:46 | Yes:Yes |
| sim3 | 4a | 1 | 4a | 1 | 854k | 99.9 | 0 | Yes |

This “multi-mapping” of reads between references in a co-infection is also evident in the plots produced by *HCVTyper* of the reference coverage and nucleotide variation in the mapping file (Figure 2 and Figure S1). Here we can clearly see from the mapping variation (panel D in both figures) that the HCV genome is most conserved in the 5’ and 3’ ends, in addition to smaller regions in the core gene, NS3 and NS5B polymerase (Fricke et al. 2015). Despite this, the consensus sequences for the two genotype 1 variants are almost identical to their respective references. For genotype 3a the multi-mapping of reads had a somewhat larger impact, resulting in 46 differences to the reference. Nevertheless, the consensus sequences were in all cases >99% identical to the reference.

**Figure 1:**
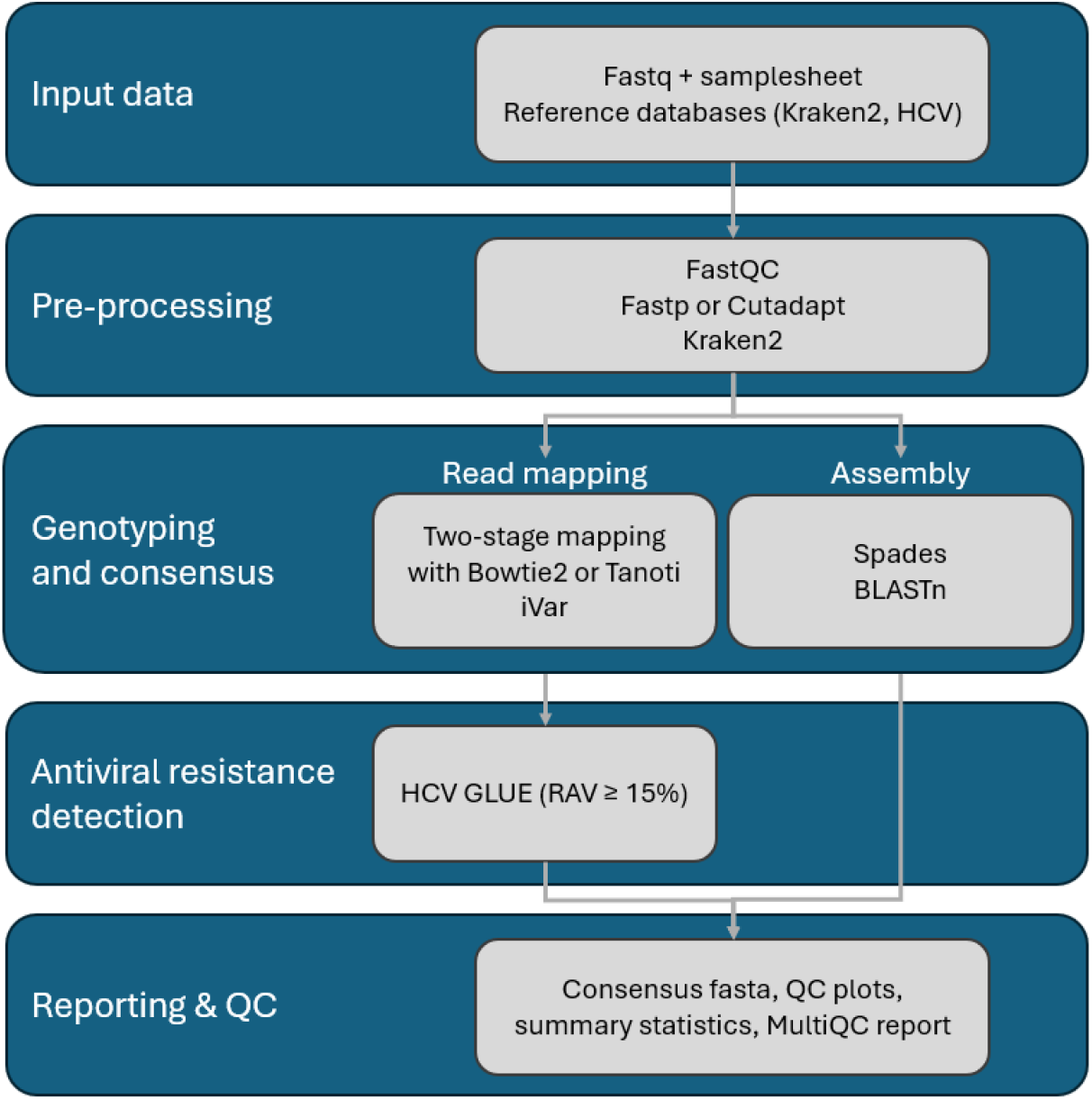
Schematic overview of the main steps in *HCVTyper*

**Figure 2:**
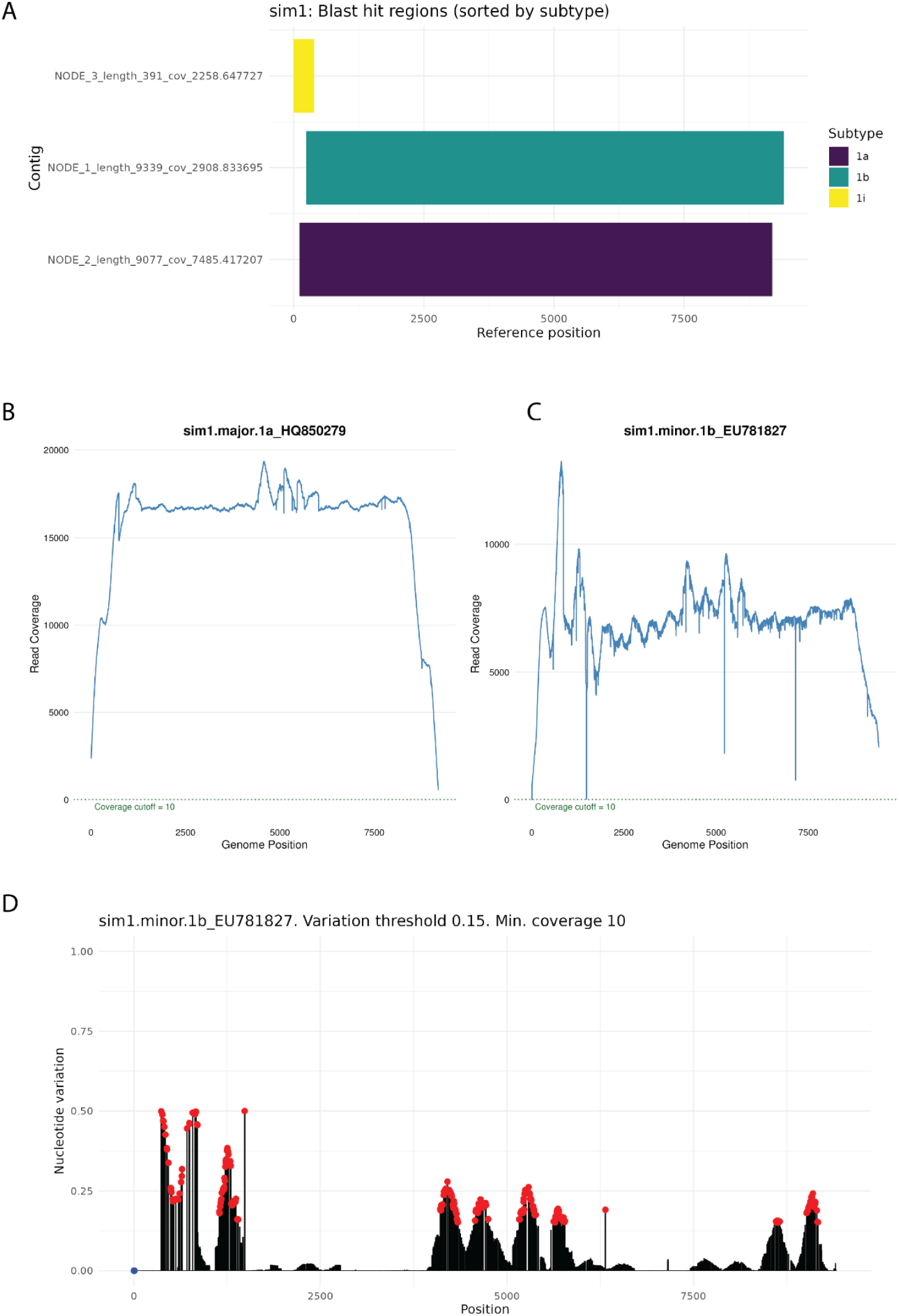
Analysis of the sim1 dataset (1a:1b co-infection). Figures that demonstrate some of the QC plots produced by the pipeline. A) Visualizations of the BLASTn search of the *de novo* assembled contigs against the HCV-reference set provided by the pipeline. The x-axis indicates the start and stop positions of the blast hits on the reference sequence. The different contigs produced by Spades are shown on the y-axis and colored according to their HCV subtype as identified by the pipeline. B) Coverage from the mapping against the identified nearest major HCV reference sequence. The x-axis indicates the genome position, the y-axis indicates the number of mapped reads (duplicates removed). The text “Coverage cutoff = 10” indicates the minimum read depth required to call a nucleotide in the consensus sequence. Regions with coverage below 10, if any present, are shaded pink. C) Same as B) but for the identified minor strain mapping. D) Visualization of the mapping variation (noise) in the mapping against the nearest minor strain reference. The x-axis indicates the genome position and the y-axis shows the degree of “Nucleotide variation” per site. This is calculated as one minus the frequency of the most common nucleotide at that position (i.e., 0.0 indicates a single nucleotide (no variation), 0.5 indicates 50/50 of two nucleotides, etc.). The blue dot highlights zero coverage regions and the red dots highlight highly variable regions (>0.15).

*de novo* assembly of the reads reconstructed full genomes as single contigs for both the 2a and 3a genomes of the sim2 dataset (Figure S2), while the sim1 dataset, which was a mixture of the more closely related genotypes 1a and 1b, resulted in an additional short contig with a BLAST hit to subtype 1i. This is likely an artifact caused by the multi-mapping of reads from the two genotypes. Such fragments are not unexpected when analyzing closely related viral genomes and do not represent a pipeline limitation, as the primary purpose of the *de novo* assemblies is to support the mapping-based genotyping.

As expected, for the simulated single-genotype dataset (sim3), the pipeline accurately identified the 4a genotype, and both the mapping and *de novo* assembly produced complete genome coverage with no spurious contigs (Figure S2).

#### “Real-world” data

We further evaluated *HCVTyper* on publicly available mixed-genotype datasets from Thomson et al. (2016) and Qiu et al. (2015) (Table 2). These datasets were obtained from Illumina capture sequencing of plasma samples from HCV-infected individuals, mixtures of plasma samples containing different genotypes, and in vitro RNA transcripts of known HCV strains mixed at defined ratios. Fastq-files were downloaded using the nf-core/fetchngs (Patel et al. 2024) pipeline by supplying the SRA accessions provided in the table. *HCVTyper* was run by setting the thresholds for minimum mapped reads (--minRead) and minimum coverage (--minCov) to 5 (see Implementation).

**Table 2:** HCVTyper run on “real-world” data. “SRA accession” refers to the accession number linking to the fastq files in the NCBI SRA database. “Theoretical genotypes” shows the expected mixtures and their relative abundances, as indicated in “Theoretical ratio”. “Identified genotypes”, “Mapped reads ratio”, “Mapped reads” and “Coverage (%)” are the same as in Table 1 except that the mapped read ratios were expressed relative to the smaller value (set to 1). Only mapped read counts greater than 1 000 are rounded to the nearest thousand. Differences between identified and theoretical genotypes are highlighted in red color. ^a^Datasets taken from (Thomson et al. 2016). ^b^Datasets taken from (Qiu et al. 2015). ^c^Approximate ratios noted in the original publication. ^d^A minimum of 10 mapped reads was required for coverage calculation; hence, zero coverage does not imply that no reads were mapped.

| SRA accession | Theoretical genotypes | Theoretical ratio | Identified genotypes | Mapped reads ratio | Mapped reads | Coverage (%) |
| --- | --- | --- | --- | --- | --- | --- |
| ERR1810511<sup>a</sup> | 1a:2a | 1:1 | 1a:2a | 1.4:1 | 8.7k:6.0k | 83.8:54.8 |
| ERR1810513<sup>a</sup> | 2a:1a | 5:1 | 2a:1a | 1.6:1 | 4.5k:2.8k | 79.8:75.0 |
| ERR1810515<sup>a</sup> | 1a:2a | 5:1 | 1a:2a | 3.6:1 | 3.2k: 0.9k | 96.8:28.3 |
| ERR1810517<sup>a</sup> | 2a:1a | 50:1 | 2a:1a | 4.4:1 | 6.1k:1.4k | 83.6:14.6 |
| ERR1810519<sup>a</sup> | 1a:2a | 50:1 | 1a:2a | 8.6:1 | 17.7k:2.0k | 99.6:33.8 |
| ERR1810521<sup>a</sup> | 2a:1a | 500:1 | 2a:1a | 4.4:1 | 9.4k:2.1k | 92.7:10.7 |
| ERR1810523<sup>a</sup> | 1a:2a | 500:1 | 1a: - | 1: - | 6.3k: - | 99.2: - |
| ERR1810525<sup>a</sup> | 2a:1a | 5000:1 | 2a: - | 1: - | 3.9k: - | 75.3: - |
| ERR1810527<sup>a</sup> | 1a:2a | 5000:1 | 1a:2a | 8.6:1 | 7.4k:864 | 99.3:11.2 |
| ERR1810497<sup>a</sup> | 1b:3a | 1:1<sup>c</sup> | 3a:1b | 2.3:1 | 84:36 | 0:2.9<sup>d</sup> |
| ERR1810499<sup>a</sup> | 3a:5a | 1:1<sup>c</sup> | 5a:3a | 1.3:1 | 23:18 | 0:0<sup>d</sup> |
| ERR1810501<sup>a</sup> | 3a:1a | 10:1 | 3a:1a | 19.1:1 | 13.7k:716 | 99.0:16.1 |
| ERR1810503<sup>a</sup> | 1a:3a | 10:1 | 1a:3a | 15.4:1 | 17.3k:1.1k | 100:11.5 |
| ERR1810505<sup>a</sup> | 1a:3a | 1:1 | 3a:1a | 1.2:1 | 8.0k:6.5k | 98.9:99.5 |
| ERR1810507<sup>a</sup> | 1b:1a | 4:1 | 1a:3a | 9.6:1 | 16.7k:1.8k | 99.8:25.5 |
| ERR1810509<sup>a</sup> | 4d:1b | 10:1 | 4d: - | 1: - | 8.5k: - | 99.3: - |
| SRR1762352<sup>b</sup> | 1b:3a | 9:1 | 1b:3a | 6.8:1 | 15.9k:23.4k | 98.7:95.7 |
| SRR1762353<sup>b</sup> | 1b:3a | 9:1 | 1b:3a | 5.6:1 | 11.4k:20.2k | 99.5:95.0 |
| SRR1762354<sup>b</sup> | 1b:3a | 1:1 | 1b:3a | 1.1:1 | 98.9k:89.9k | 99.2:99.2 |
| SRR1762355<sup>b</sup> | 1b:3a | 1:1 | 1b:3a | 1.5:1 | 12.5k:84.9k | 99.0:97.9 |

For the majority of samples, the pipeline correctly identified both genotypes in the co-infection datasets. Performance remained robust across a wide range of input ratios, including cases where the minor genotype contributed only ∼2-10% of reads. As expected, detection of minority strains declined in samples with extreme input ratios (500:1 or 1:5000). In these cases, the minor strain was not reported because the number of mapped reads fell below the thresholds for mapped reads and coverage. For example, in ERR1810523 (500:1 mixture), only genotype 1a was reported, while in ERR1810525 (1:5000 mixture), only genotype 2a was recovered. One notable exception was ERR1810509, a 10:1 mixture of 4d and 1b, in which the minor genotype (1b) was not detected despite the moderate difference in expected abundance. This sample contained relatively few total reads, and it is not uncommon that minority strains do not reach sufficient number of mapped reads for confident detection.

Only a single misclassification was observed: ERR1810507 (true mixture 1a:1b) was reported as 1a:3a. Closer inspection however, revealed that the pipeline had actually recovered *de novo* assembled fragments of a complete 1b genome (Figure S3). It also identified 791 reads mapping to 1b reference genomes in the first mapping (Supplementary file 1), with up to 11.7% coverage (calculated from Supplementary file 2). However, reference sequences belonging to genotype 3a had more reads mapped to them, and higher coverage (Figure S4). This strongly suggests contamination with genotype 3a in this sample and shows the need for manual inspection of QC metrics and assemblies in clinically relevant samples.

The differences in mapped reads between the major and minor strains, did not accurately reflect the theoretical expectations across samples. This discrepancy likely reflects common biases in NGS data, particularly when capture-based sequencing and multiple amplification steps during library preparation are involved (Aird et al. 2011). In addition, the libraries analyzed here were dominated by human reads (Qiu et al. 2015; Thomson et al. 2016), which is common when working with human samples. This will further exacerbate amplification bias and distort the true viral diversity. And importantly,

*HCVTyper* reports mapped reads after removal of duplicate reads, which often constitute a substantial portion of the total reads. Consequently, the reported read counts do not directly reflect total sequencing depth or true input ratios. Despite this, *HCVTyper* consistently distinguished between the major and minor genotypes present in each sample.

Together, these results demonstrate that *HCVTyper* reliably detects co-infections across a wide dynamic range, but interpretation of borderline cases benefits from careful review of the provided mapping statistics, coverage profiles, and QC reports.

### Pipeline performance

On an 8 CPU node Linux computer with 50 GB RAM, a single sample (1 million Illumina read pairs) completes in about 20 minutes (depending on whether there is a single or co-infection). When running all processes as Docker containers, some time will be used to download and stage the different container images. Also, if downloading the default Kraken2 database, downloading and unpacking will also consume a significant amount of time. We therefore recommend downloading and storing a local copy of the Kraken2 database if the pipeline is frequently used.

The strength of using Nextflow to orchestrate the pipeline is that it efficiently scales the execution when processing multiple samples. A typical scenario is to run a batch of samples analyzed together on a high-throughput sequencing platform. To evaluate such typical performance, we ran a batch of 20 samples from an Illumina MiSeq sequencer on a 16 CPU node Linux machine, with 230 GB RAM (Figure 3). The pipeline completed in 1h14min, with the most time-consuming process being HCV-GLUE and the most memory intensive process being Kraken2. However, Kraken2 which was run against the PlusPFP-8 database, did not consume more than 10 GB RAM. The HCV-GLUE process is run only once on all samples together, whereas the other processes run independently for each sample in parallel; therefore, the timing of HCV-GLUE needs to be divided by the number of samples. Overall, these resource requirements are well within capacity of computing environments typically available in most laboratories.

**Figure 3:**
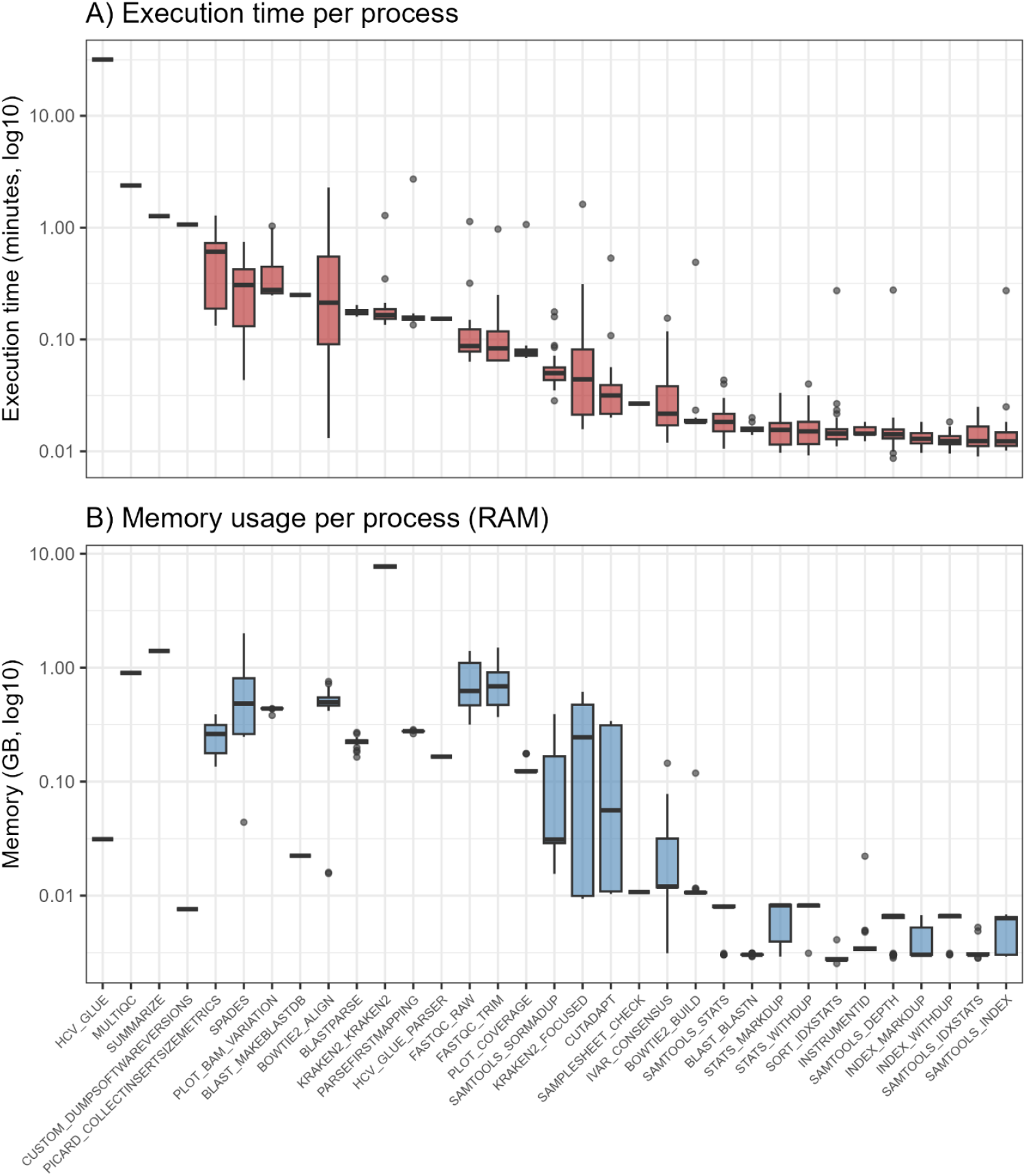
Execution time and memory usage per pipeline process. Time elapsed (A) and memory usage (B) in an analysis of 20 samples of Illumina MiSeq paired-end reads run on a Linux computer with 16 CPUs and 230GB RAM available. The values for all samples for each process are summarized as boxplots with the median shown as the thick line, the interquartile range (75^th^-25^th^ percentile) shown in the boxes, the whiskers showing the lowest and highest data points, and the outliers displayed as black dots.

Another strength of using Nextflow is that the pipeline can be stopped and re-started from where it left off, as well as adding samples to a stopped session, and the pipeline will add these and pick up the others where it left off.

By default, the pipeline will store output files from most intermediate steps (e.g. BAMs, alignment stats, log files) as these can be useful for auditing. However, as these may take up unnecessary disk space, it is possible to customize each step of the pipeline to control which intermediate files will be stored.

## Discussion

HCVTyper provides an end-to-end, containerized workflow for the assembly, genotyping, co-infection detection, and resistance analysis of HCV genomes from Illumina sequencing data. By leveraging the Nextflow and nf-core frameworks, the pipeline is portable, scalable, and fully reproducible across computing environments. Compared to more general virus assembly pipelines, *HCVTyper* offers a tailored approach that accounts for HCV’s high genetic variability and clinical relevance of resistance-associated variants.

A central design principle of *HCVTyper* is flexibility. Rather than applying fixed thresholds to define major and minor strains, the pipeline provides alignment metrics, coverage statistics, and QC reports, allowing users to set criteria appropriate for their study. This is particularly important in co-infected samples, where conserved genomic regions can lead to reads mapping across multiple genotypes. This can inflate apparent support for minor strains and introduce errors into consensus sequences, particularly for closely related subtypes. In controlled mixtures where the true diversity is known, *HCVTyper* can often detect minor variants even when supported by relatively few reads. However, in routine diagnostics or surveillance samples, particularly when capture-based enrichment is used, background noise and non-HCV reads can generate spurious low-level signals. Based on our experience, a minimum of ∼30% genome coverage and ≥500 mapped reads provides a practical balance between sensitivity and specificity, though stricter or looser thresholds may be appropriate depending on the application.

An important point is that the number of mapped reads does not necessarily reflect the true abundance of viral strains in a sample. In the simulated datasets, the mapped read ratios closely matched the expected proportions, which is not surprising since the data contained only HCV references, and we did not simulate PCR-or capture-related artifacts. In contrast, the real-world datasets were much more complex. Most reads in these samples were of human origin (Qiu et al. 2015; Thomson et al. 2016), which is typical when sequencing viruses directly from human samples. In addition, enrichment biases and amplification steps during library preparation can strongly influence the observed read ratios. Thus, while mapped read counts may provide a rough indication, they cannot be used to accurately reconstruct viral abundances from real-world data. The aim of *HCVTyper* is therefore not to quantify strain abundance, but to robustly identify the genotypes or strains present in each sample.

Another complication arises from the existence of naturally occurring recombinant HCV strains, in which sequences from distinct genotypes are combined within the same viral genome (Hedskog et al. 2015). Such recombinants can be difficult to distinguish from co-infections, since reads may map to different parental genotypes depending on the genomic region considered. A well-described example is the 2k/1b recombinant (Raghwani et al. 2012; Zakalashvili et al. 2018). To account for this, the pipeline is designed not to report simultaneous co-infections involving genotype 2 and genotype 1 if a 2k/1b recombinant is identified. This reduces the risk of misinterpreting a true recombinant as a mixed infection.

For reference-based assembly, *HCVTyper* implements a two-step mapping strategy, similar to the approach taken in the MiCall pipeline. Reads are first aligned to a broad set of references spanning all HCV genotypes, followed by re-mapping to the most similar candidate(s). This reduces the risk of bias from an inappropriate initial reference choice, a common challenge in viral genome analysis. *HCVTyper* will also perform *de novo* assembly of the reads and genotyping of the resulting contigs. However, *de novo assembly*, especially on capture or amplicon data, often produce highly fragmented assemblies that can be difficult to piece together. Therefore, reference-based mapping remains the backbone of consensus sequence generation and RAV identification.

Among the available mapping tools, Bowtie2 is the default due to its speed, low disk usage, and active maintenance. Tanoti, while historically popular in HCV research (Thomson et al. 2016), is slower, more resource intensive, and no longer actively maintained, and is therefore included only as an alternative.

It is also important to note that *HCVTyper* is not a haplotype reconstruction tool. The consensus sequences it produces should be interpreted as representative of the predominant viral strains present, rather than full haplotype-resolved quasispecies.

For antiviral resistance detection, *HCVTyper* integrates HCV-GLUE. A key advantage of this approach is that RAVs are identified directly from the mapping (BAM) files, which allow detection of minority variants within the viral population. By default, *HCVTyper* reports RAVs present at frequencies as low as 15%, but this threshold can be adjusted by the user to suit different research or clinical contexts and also control the balance between sensitivity and specificity. In addition to resistance detection, HCV-GLUE provides robust phylogenetic-based genotyping. This can be compared against the mapping-based genotyping by *HCVTyper* as an additional control. At present, HCV-GLUE is only supported in *HCVTyper* when the pipeline is run with Docker, which may limit use in some environments. Future development of Singularity- or Conda-compatible builds would improve accessibility, especially on high-performance computing clusters where Docker is not available.

Overall, *HCVTyper* complements existing virus analysis pipelines by providing a portable, open, and HCV-specific workflow that is suitable for both research and public health contexts. Its modular design allows integration of updated reference sets, resistance annotation databases, and future methodological improvements. By leveraging nf-core community standards, the pipeline will remain maintainable and extensible as sequencing technologies and clinical requirements evolve. Although the pipeline is specifically designed for analyzing data from Illumina capture sequencing, it is also able to assemble genomes from shot-gun sequencing or tiling PCR sequencing, given that primer sequences are removed from the reads prior to analysis. In principle, the workflow could also be applied to other viral agents by replacing the reference dataset, although in such cases HCV-GLUE would not be available. We invite users and developers to try the pipeline on their own HCV data and contribute improvements via the GitHub repository.

## Acknowledgements

Thanks to the nf-core community on Slack for providing help and support.

