## Supplementary material for "HCVTyper: A Nextflow pipeline for hepatitis C virus genome assembly, genotyping and antiviral resistance detection"

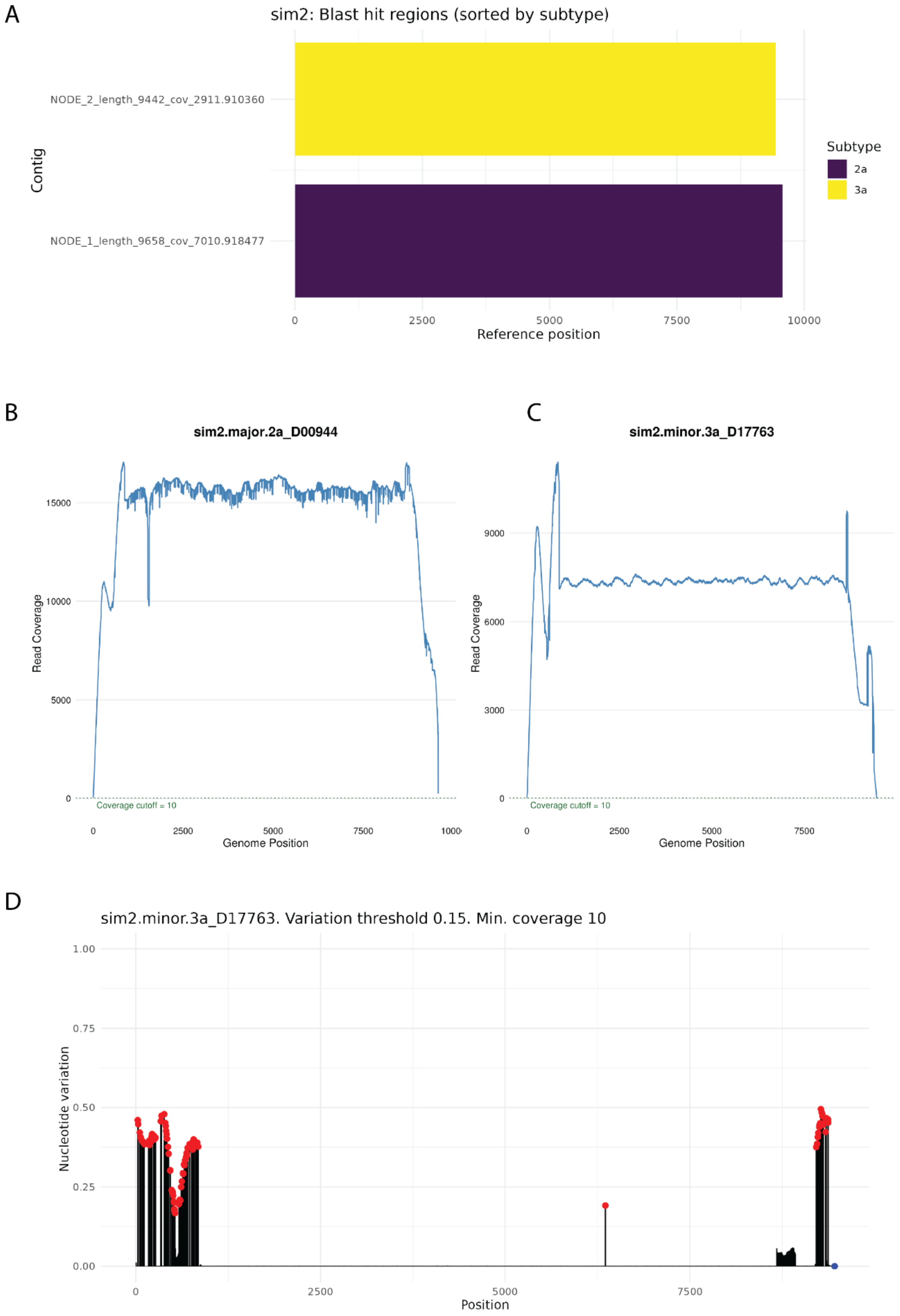

**Figure S1: Analysis of the sim2 dataset (2a:3a co-infection)**. Figure legend same as Figure 2.

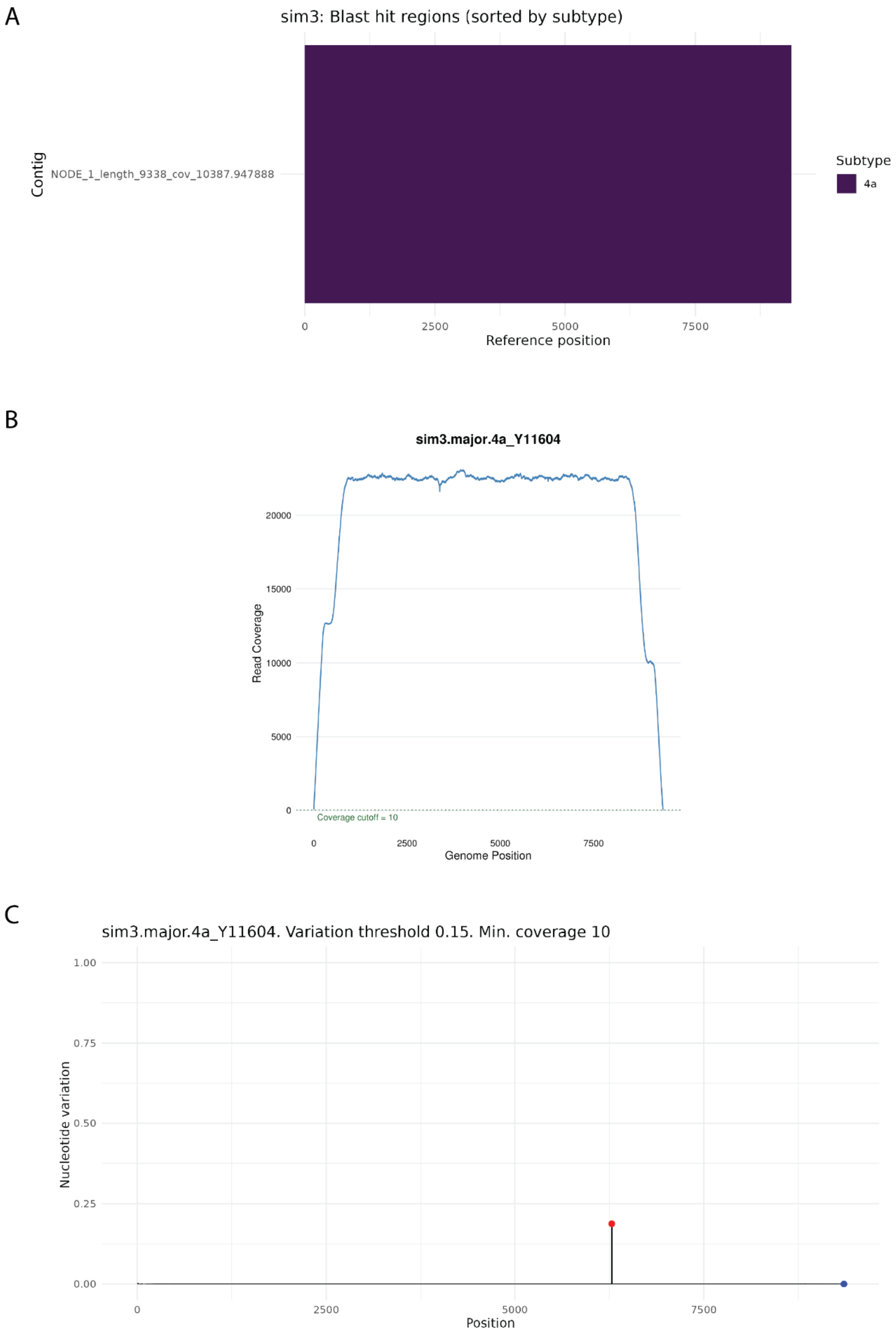


**Figure S2: Analysis of the sim3 dataset (4a infection).** Figure legend same as Figure 2. Note that there are no plots for any “minor” strain as only a single genotype was simulated for this sample.

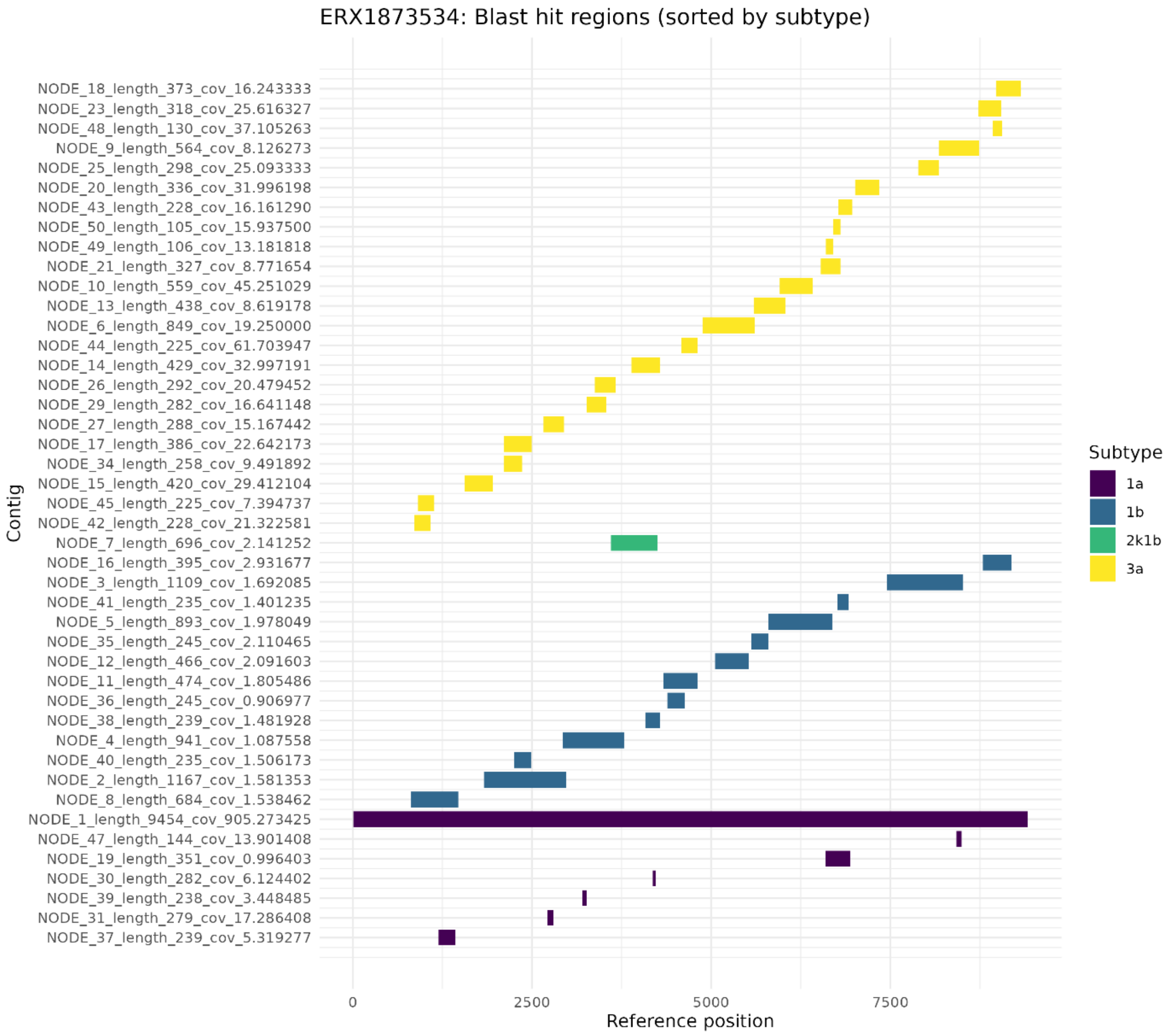


**Figure S3. Genotyping of *de novo* assembled contigs from sample ERR1810507.** See Figure 2 legend for description of figure. Notice that three different genotypes are detected, 1a, 1b and 3a. The short fragment assigned to 2k1b is most likely an artifact because this is a recombinant reference with genomic similarity to 1b.

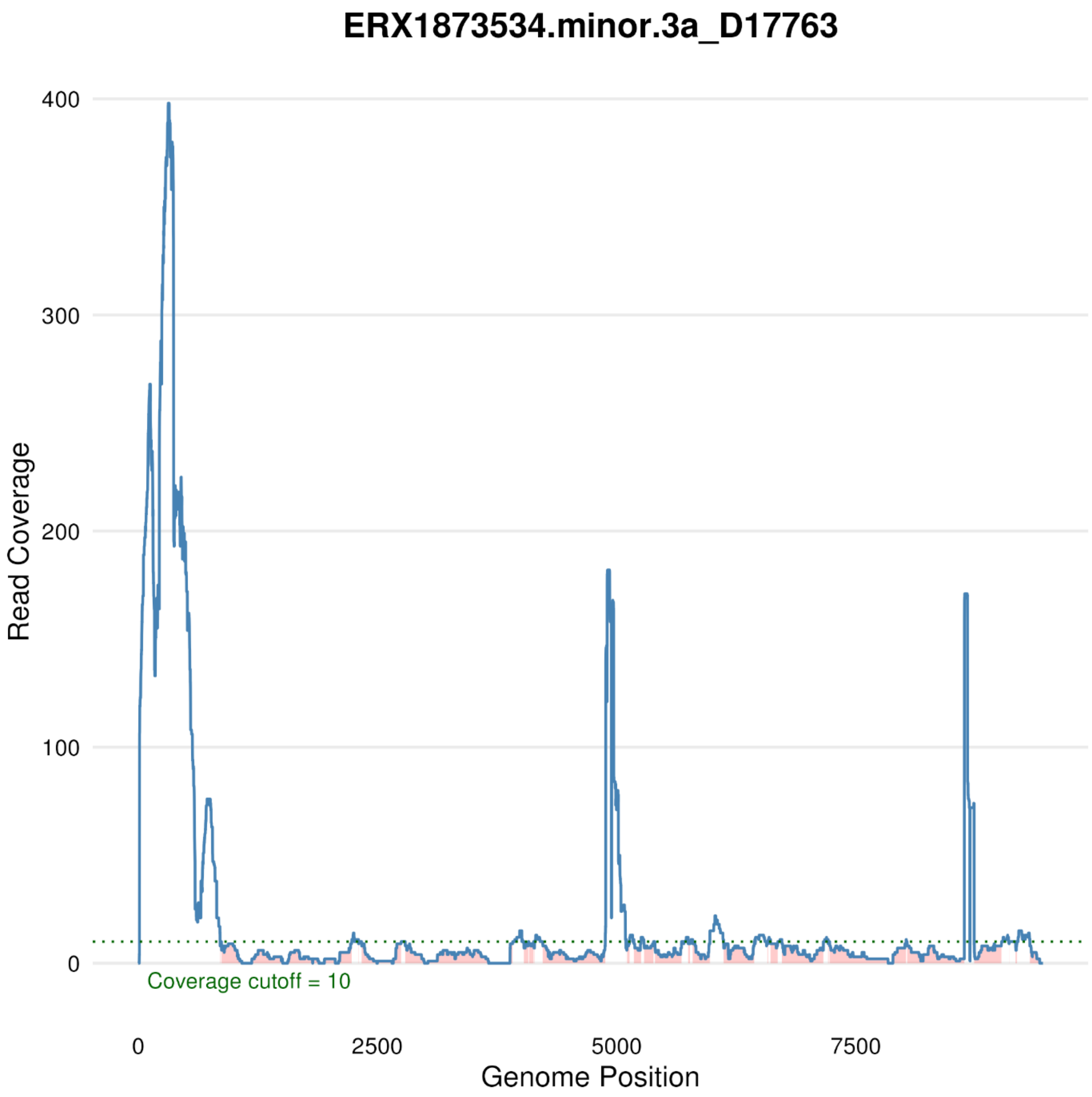


**Figure S4. Coverage from the mapping against the selected minor reference from sample ERR1810507.** See Figure 2 legend for description of figure.

**Supplementary file 1.** Output from *samtools idxstats* run on the bam-file from the first mapping against all HCV references, with duplicates included. The file is sorted according to the number of mapped reads (column 3).

**Supplementary file 2.** Output from *samtools depth* run on the bam-file from the first mapping against all HCV references, with duplicates included. The coverage of each reference was calculated by dividing the sum of each position with a coverage greater than four by the total number of positions.
